## Supplementary_information for "Developmental stage shapes the realized energy landscape for a flight specialist"

#### This PDF file includes:

Figure S1

Figure S2

Legend for Video S1

Table S1

#### Other supplementary information for this paper include the following:

Video S1

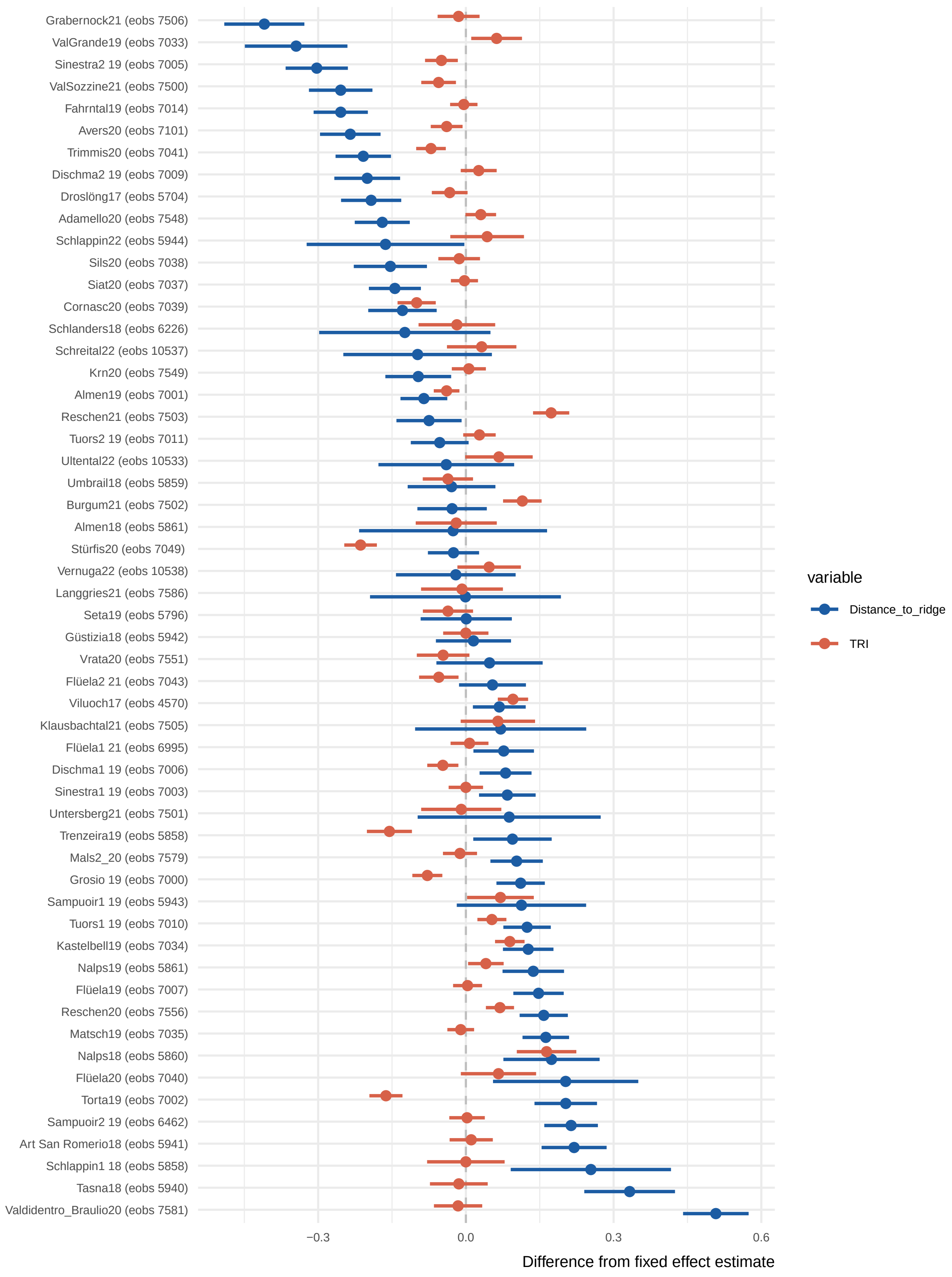


Figure S1: Individual-specific slopes for Topographic Ruggedness Index (TRI) and distance to ridge line. The difference between each individual's estimate from the fixed effect estimate (Fig. 1) is shown. Individuals are ordered based on their distance to ridge line estimates.

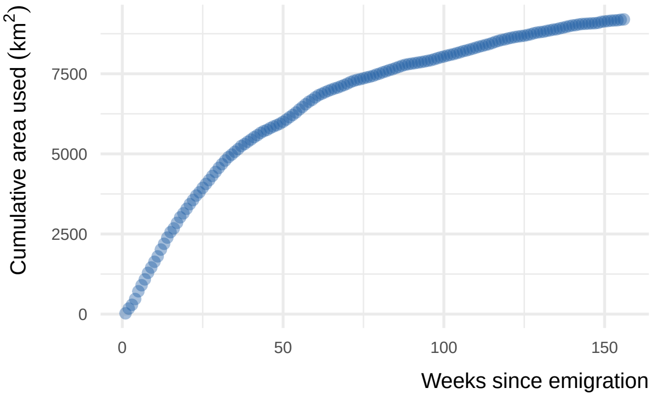


Figure S2: Cumulative area used by juvenile golden eagles during each week after emigration. Used areas were calculated by extracting the commuting flights for each week, converting these to line objects, overlapping the lines with a raster of 100*100 km cell size, counting the number of overlapping cells and calculating the area that they covered. The predicted flyable area for juvenile golden eagles in the Alpine region for the same period of time is shown in Fig. 4.

### Legend for Video S1

The realized energy landscape of the aging golden eagles. We used a step-selection approach to determine how juvenile golden eagles responded to topographic conditions during their commuting flights. Topography is a predictor of uplift potential and can be used as a proxy for energy availability for soaring birds. We used a step-selection model to predict flyability across the Alpine region from one week to three years after emigration. The fundamental energy landscape, defined as the total amount of energy available in the landscape, is constant, but the realized energy landscape, here estimated as flyability, changes. This is because the birds' ability to perceive and exploit the energy within the landscape improves as they age, making the landscape cheaper to traverse. Flyability quantifies the suitability of a location for efficient flight, with higher values indicating areas where the bird is more likely to benefit from favorable uplifts. It represents the realized energy landscape that a bird can exploit based on the given topographic conditions and its own cognitive and locomotor abilities.

Table S1. Details of bio-logging data included in the study. The country and year of tagging for each individual, and the tracking duration (in terms of weeks since dispersal) that each individual contributed to the analysis, are reported. All individuals carried Bird Solar Tags manufactured by e-obs GmbH, Germany (either 45 gr or 25 gr devices). Individual local identifiers match those included in Movebank studies "LifeTrack Golden Eagle Alps" and "LifeTrack Golden Eagle Alps Public".

| Individual local identifier | Country | Tagging year | Weeks post-dispersal |
|---|---|---|---|
| Adamello20 (eobs 7548) | Italy | 2020 | 106 |
| Almen18 (eobs 5861) | Switzerland | 2018 | 1 |
| Almen19 (eobs 7001) | Switzerland | 2019 | 156 |
| Art San Romerio18 (eobs 5941) | Switzerland | 2018 | 60 |
| Avers20 (eobs 7101) | Switzerland | 2020 | 105 |
| Burgum21 (eobs 7502) | Italy | 2021 | 42 |
| Cornasc20 (eobs 7039) | Switzerland | 2020 | 59 |
| Dischma1 19 (eobs 7006) | Switzerland | 2019 | 156 |
| Dischma2 19 (eobs 7009) | Switzerland | 2019 | 83 |
| Droslöng17 (eobs 5704) | Switzerland | 2017 | 102 |
| Fahrntal19 (eobs 7014) | Italy | 2019 | 156 |
| Flüela1 21 (eobs 6995) | Switzerland | 2021 | 61 |
| Flüela19 (eobs 7007) | Switzerland | 2019 | 156 |
| Flüela2 21 (eobs 7043) | Switzerland | 2021 | 49 |
| Flüela20 (eobs 7040) | Switzerland | 2020 | 8 |
| Grabernock21 (eobs 7506) | Italy | 2021 | 61 |
| Grosio 19 (eobs 7000) | Italy | 2019 | 156 |
| Güstizia18 (eobs 5942) | Switzerland | 2018 | 83 |
| Kastelbell19 (eobs 7034) | Italy | 2019 | 141 |
| Klausbachtal21 (eobs 7505) | Germany | 2021 | 2 |
| Krn20 (eobs 7549) | Slovenia | 2020 | 80 |
| Langgries21 (eobs 7586) | Austria | 2021 | 46 |
| Mals2_20 (eobs 7579) | Italy | 2020 | 65 |
| Matsch19 (eobs 7035) | Italy | 2019 | 156 |
| Nalps18 (eobs 5860) | Switzerland | 2018 | 42 |
| Nalps19 (eobs 5861) | Switzerland | 2019 | 60 |
| Reschen20 (eobs 7556) | Italy | 2020 | 130 |
| Reschen21 (eobs 7503) | Italy | 2021 | 60 |
| Sampuoir1 19 (eobs 5943) | Switzerland | 2019 | 9 |
| Sampuoir2 19 (eobs 6462) | Switzerland | 2019 | 72 |
| Schlanders18 (eobs 6226) | Italy | 2018 | 4 |
| Schlappin1 18 (eobs 5858) | Italy | 2018 | 3 |
| Schlappin22 (eobs 5944) | Switzerland | 2022 | 7 |
| Schreital22 (eobs 10537) | Germany | 2022 | 19 |
| Seta19 (eobs 5796) | Switzerland | 2019 | 19 |
| Siat20 (eobs 7037) | Switzerland | 2020 | 107 |
| Sils20 (eobs 7038) | Switzerland | 2020 | 62 |
| Sinestra1 19 (eobs 7003) | Switzerland | 2019 | 156 |
| Sinestra2 19 (eobs 7005) | Switzerland | 2019 | 156 |
| Stürfis20 (eobs 7049) | Switzerland | 2020 | 100 |
| Tasna18 (eobs 5940) | Switzerland | 2018 | 17 |
| Torta19 (eobs 7002) | Switzerland | 2019 | 69 |
| Trenzeira19 (eobs 5858) | Italy | 2019 | 29 |
| Trimmis20 (eobs 7041) | Switzerland | 2020 | 119 |
| Tuors1 19 (eobs 7010) | Switzerland | 2019 | 156 |
| Tuors2 19 (eobs 7011) | Switzerland | 2019 | 156 |
| Ultental22 (eobs 10533) | Italy | 2022 | 6 |
| Umbrail18 (eobs 5859) | Switzerland | 2018 | 37 |
| Untersberg21 (eobs 7501) | Germany | 2021 | 2 |
| Valdidentro_Braulio20 (eobs 7581) | Italy | 2020 | 24 |
| ValGrande19 (eobs 7033) | Italy | 2019 | 46 |
| ValSozzine21 (eobs 7500) | Italy | 2021 | 60 |
| Vernuga22 (eobs 10538) | Italy | 2022 | 12 |
| Viluoch17 (eobs 4570) | Switzerland | 2017 | 137 |
| Vrata20 (eobs 7551) | Slovenia | 2020 | 16 |
